# Functional Redundancy Permits Enzyme Specialisation Without Gene Duplication: A Minimal Model

**DOI:** 10.64898/2026.10.02.756335

**Authors:** Alp Duygu, Haziq Moinudeen

## Abstract

Gene duplication is the standard explanation for how multifunctional enzymes specialise: a spare copy is freed from maintaining the ancestral functions, bypassing functional entrenchment. Yet proteins often share overlapping weak activities, so other contributors may already buffer the loss of one. We ask whether this overlap alone permits specialisation without duplication, using a minimal origin–fixation model in which weak generalist proteins jointly supply 120 organismal reactions. Fitness depends on aggregate reaction performance with diminishing returns, and pleiotropic mutations alter activity profiles under structural incompatibility. Protein number is fixed and fitness has no reward for narrow profiles. At five fixed substitutions per protein, effective breadth falls across redundancies of 5 to 15 proteins per reaction. Under a loss-biased mutation kernel, greater redundancy generally produces stronger specialisation while better preserving the least-supplied reaction, consistent with an analytical result that redundancy lowers the selective cost of losing one contribution. A symmetric kernel produces stronger specialisation still, so loss bias is not required, although the kernels differ in magnitude distribution. Distributed overlap therefore lets existing proteins specialise without losing reaction coverage. Duplication remains a major route, but this model shows it is not a necessary first step.

## I. INTRODUCTION

Enzymes organise cellular chemistry by accelerating particular reactions, but specificity does not imply an absolute restriction to one substrate or one catalytic transformation. Many enzymes also possess secondary activities that are weak under their usual physiological conditions yet can provide starting points for evolutionary change [1, 2]. Reconstructed ancestral enzymes offer concrete examples of broader catalytic repertoires preceding the divergence of specialised descendants. In fungal sugar metabolism, for example, ancestral reconstruction and biochemical measurements traced the separation of maltase and isomaltase activities from a promiscuous precursor [3]. Such cases pose a general problem: how can a protein become more specialised when reducing its other activities may compromise functions that the organism still requires?

Gene duplication provides a well-established answer. In Ohno’s classical account, an additional gene copy can relax the requirement that every copy maintain the ancestral function, allowing functional divergence [4]. Neofunctionalisation describes the acquisition of a new function, whereas subfunctionalisation describes the partitioning of ancestral functions among copies. The duplication– degeneration–complementation model showed how complementary losses, particularly in regulatory subfunctions, can favour retention of both duplicates [5]. These routes need not be mutually exclusive in enzyme evolution: changes in dosage, catalytic specificity, and physiological role can occur along the same lineage [3, 6]. Duplication is therefore both an empirically important process and a useful way to separate competing functional demands on a multifunctional protein.

The order of innovation and duplication is also consequential. In the innovation–amplification–divergence model, an initially weak activity first becomes useful, selection favours increased gene dosage, and subsequent mutations improve the activities of different copies [7]. Näsvall and colleagues observed this process experimentally in a bacterial system requiring a bifunctional enzyme to support two biosynthetic pathways [8]. This result demonstrates that duplicate retention need not depend on an extended period of neutral exploration. More generally, duplication-based explanations encompass several orders of functional change and selection, rather than a single obligatory sequence of events [6].

Enzyme recruitment supplies the biochemical connection between weak secondary activity and a new physiological role. Jensen proposed that broad substrate specificities could allow existing enzymes to be recruited into additional metabolic functions, with subsequent duplication and divergence permitting greater specialisation [9]. Work on catalytic promiscuity developed this idea by identifying how an active site adapted to one reaction can also catalyse other transformations [1, 2]. Promiscuous activities are not necessarily fixed by the constraints on the principal activity. Directed-evolution experiments found that secondary activities can change substantially while the native activity changes much less [10]. Over longer adaptive trajectories, however, improvement can encounter diminishing returns and asymmetric trade-offs between activities [11]. These observations motivate models that distinguish the distribution of mutational effects from the selective value of the resulting phenotype.

The phenotype in which an enzyme operates extends beyond that enzyme’s own activity profile. Its physiological contribution depends on the other proteins present, the metabolic network, and the environment, as emphasised in systems-level accounts of enzyme evolution [12]. Experimental multicopy-suppression screens also show that overexpression of a noncognate gene can rescue some metabolic deficiencies [13]. Such rescue can involve alternative reactions or pathways as well as overlapping activities; it does not establish that all catalytic contributions are interchangeable. It nevertheless illustrates why the effect of modifying one protein cannot generally be inferred from its isolated catalytic profile. Functional overlap is a property of the surrounding biochemical system, and need not be confined to a recently duplicated pair of genes.

This perspective suggests a narrower theoretical question. Suppose several weak generalists contribute to each required reaction and selection acts on their aggregate catalytic capacity. Can the existing proteins become more specialised while that shared reaction set remains supplied? A loss of activity in one contributor may be buffered by the others, whereas a gain may have little value if a reaction is already well supplied. Both effects depend on the distribution of activity across the proteome. We examine this question at fixed protein number within each evolutionary trajectory. This choice sets aside the origin of the ancestral repertoire, which may itself have involved duplication or other processes, and isolates the subsequent reorganisation of existing catalytic contributions. It also separates specialisation within an established reaction set from the evolution of new organismal chemistry.

Here we construct a minimal stochastic model in which organismal fitness depends on the combined catalytic performance of a set of required reactions. Proteins begin as heterogeneous weak generalists with overlapping activity profiles. Mutations alter several activities within coarsegrained structural loops, while a local incompatibility constraint limits the combinations of activities that can be reached. There is no direct fitness reward for protein-level specialisation and no gene duplication during the simulated transition.

We vary the initial redundancy of the proteome, the mutation kernel, and structural incompatibility. Because increasing redundancy also increases the number of proteins in the model, evolutionary outcomes are compared at equal numbers of fixed substitutions per protein rather than at equal total substitution counts. Under this comparison, increasing redundancy generally produces greater narrowing of protein activity profiles under loss-biased mutation, while the least well supplied organismal reactions become better buffered. The symmetric mutation kernel produces stronger specialisation overall, indicating that biochemical loss bias is not required for the effect. The model therefore identifies a route by which distributed functional overlap can permit existing proteins to divide catalytic labour without an additional duplication event.

## II. METHODS

### A. Proteome-level activity and fitness

We consider a proteome of *N* protein-coding loci contributing to *M* organismal reactions. Protein *i* has a non-negative activity vector **a**_*i*_ = (*a*_*i*1_, … , *a*_*iM*_), and the complete proteome is represented by the activity matrix

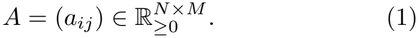

The variables *a*_*ij*_ are phenomenological catalytic activities rather than microscopic kinetic constants. Simulations use equal protein abundance, so the aggregate catalytic capacity available to reaction *j* is

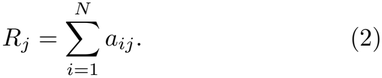

Reaction performance increases with aggregate capacity but has diminishing returns,

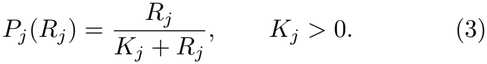

Organismal fitness is the weighted geometric mean of reaction performances,

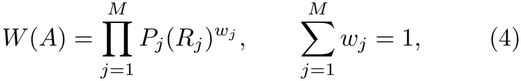

with equal weights *w*_*j*_ = 1*/M* in the simulations. If any required reaction has zero aggregate activity, fitness is zero. Equivalently,

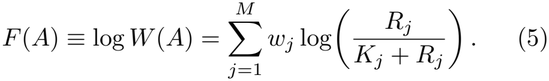

Thus the selective effect of changing one protein is determined by the reaction capacity already supplied by the rest of the proteome. The marginal value of additional activity in reaction *j* is

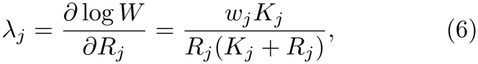

which decreases as *R*_*j*_ increases.

### B. Loop-local pleiotropic mutation and structural feasibility

Mutations act on coarse-grained structural regions, termed loops and indexed by *l*. The contribution of loop *l* on protein *i* to reaction *j* is *L*_*ilj*_ ≥ 0, with

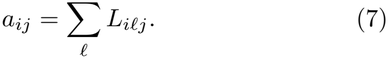

Each loop has a latent mutational support *T*_*il*_, the set of reactions whose activity can be altered by ordinary mutations of that loop. Activities that reach zero remain in *T*_*il*_ and can therefore be restored by later positive mutations; zero activity is not an absorbing state.

To represent local incompatibility among simultaneously strong activities, each loop is assigned the phenomenological structural cost

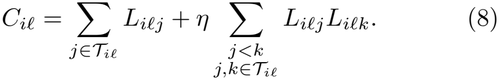

A proposed state is feasible only when

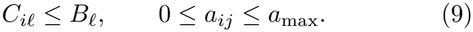

The structural term does not enter organismal fitness directly; it restricts the accessible biochemical state space. We report incompatibility using the dimensionless quantity

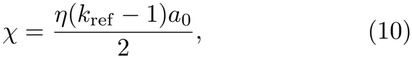

where *a*_0_ is the mean ancestral activity of a supported protein–reaction pair and *k*_ref_ is the mean ancestral loop support size. The sweep uses *χ* ∈ {0, 0.1, 0.325, 1}. For the reference architecture, *k*_ref_ = 4.25 and *a*_0_ = 0.02, so *χ* = 0.325 corresponds to *η* = 10. The limit *χ* = 0 removes the pairwise incompatibility term.

### C. Mutation kernels and support expansion

An ordinary mutation acts on one loop of one protein. Conditional on the latent support *T*_*il*_, each affected coordinate independently receives a sign *σ*_*j*_ ∈ {−1, 0, +1} .

In the loss-biased reference kernel,

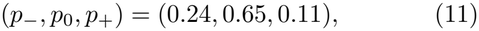

where *p*_*σ*_ = Pr(*σ*_*j*_ = *σ*). Positive and negative magnitudes are drawn from

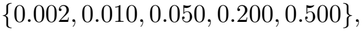

with respective probabilities

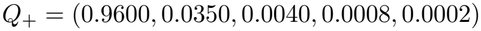

and

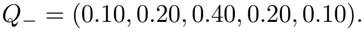

The realised activity change is

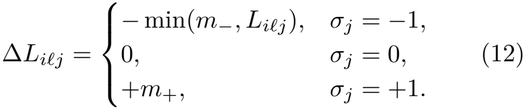

The symmetric control retains *p*_0_ = 0.65, sets *p*_−_ = *p*_+_ = 0.175, and uses the same magnitude distribution for gains and losses. The common magnitude probabilities are the elementwise mean of the reference positive and negative distributions,

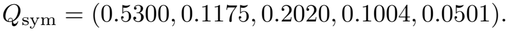

The symmetry refers to proposed signs and magnitudes; truncation at zero and feasibility filtering still affect realised changes. All other components of the model are held fixed between mutation kernels.

Rare support-expansion events occur with probability *ρ*_exp_ = 10^−3^ among mutation origins. An expansion adds a weak activity toward a reaction not currently contained in the loop’s latent support, subject to the same structural feasibility rule, and permanently adds that reaction to *T*_*il*_. The organismal reaction set *M* itself remains fixed.

### D. Ancestral generalists and redundancy sweep

All simulations use *M* = 120 organismal reactions, ancestral protein breadth *k*_*p*_ = 15, four loops per protein with support sizes (5, 4, 4, 4), initial total protein activity *A*_*i*,0_ = 0.30, *B*_*l*_ = 1, and *a*_max_ = 1.

The ancestral protein–reaction network is randomised while preserving protein-side degree and balanced reactionside degree. Within each protein, positive Gamma-distributed weights with shape parameter *α* = 8 are drawn over its 15 supported reactions *S*_*i*_ and normalised to the fixed total activity (the common Gamma scale cancels),

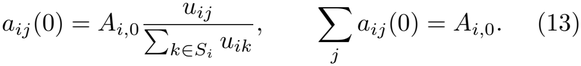

This produces weak heterogeneous generalists without placing every ancestor at the maximum of the effective-breadth statistic.

Initial mean reaction redundancy is

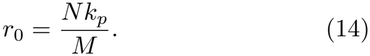

We use

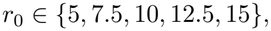

corresponding to *N* ∈ {40, 60, 80, 100, 120 ,} while *M* and *k*_*p*_ remain fixed. For every condition we set

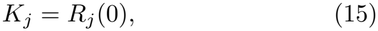

so that all reactions begin with *P*_*j*_(0) = 1*/*2, independent of redundancy. Each *K*_*j*_ is then held fixed during evolution.

### E. Origin–fixation dynamics and numerical implementation

Finite-effect mutations are evaluated using the exact selection coefficient

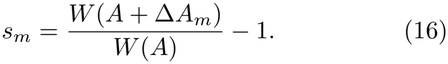

Evolution follows an origin–fixation process: each sampled substitution is fixed before the next is considered. We use the diffusion-approximation fixation probability [14], with effective population size *N*_*e*_,

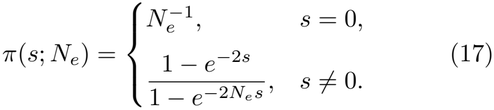

For mutational event *m* with origin rate *v*_*m*_, the fixation hazard is *h*_*m*_ = *v*_*m*_*π*(*s*_*m*_; *N*_*e*_). The total hazard *H* = ∑_*m*_ *h*_*m*_ determines the waiting time *τ∼*Exp(*H*), and the next substitution is sampled with probability *h*_*m*_*/H*. Simulations use *N*_*e*_ = 10^6^ and loop-level mutation rate *U*_loop_ = 10^−8^. Mutation catalogues are enumerated exactly for each loop support size, cached, and evaluated in vectorised form. After each fixed substitution, fixation hazards are recomputed only for loops whose mutational or fitness context can have changed. Independent parameter combinations were run in parallel.

The complete experiment is a 5 × 2 × 4 grid of redundancy, mutation kernel, and structural incompatibility, with independent stochastic replicates for each parameter combination. At each redundancy, ancestral seeds were matched across mutation kernels and incompatibility values, while evolutionary seeds differed between runs.

Initial redundancy is varied by changing protein number *N*. A fixed number of substitutions across all conditions would therefore expose each protein to fewer evolutionary events in larger proteomes. To make redundancy conditions comparable, we define evolutionary exposure as

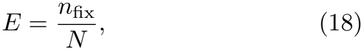

where *n*_fix_ is the total number of fixed substitutions in the proteome. Primary endpoint comparisons are made at *E* = 5 fixed substitutions per protein. The corresponding stopping points are therefore 200, 300, 400, 500, and 600 fixed substitutions for *N* = 40, 60, 80, 100, and 120, respectively. This normalisation holds the average evolutionary opportunity per protein constant while allowing the total mutation supply of the proteome to scale naturally with protein number.

### F. Summary statistics

For protein *i*, the normalised catalytic profile is

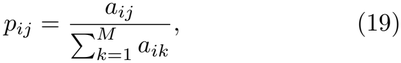

and effective functional breadth is

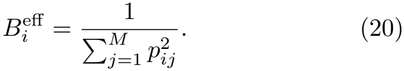

For reaction *j*, contribution shares are

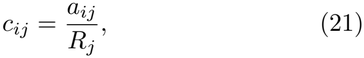

giving effective reaction redundancy

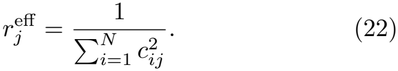

We additionally record each protein’s strongest-function share, *q*_*i*,max_ = max_*j*_ *p*_*ij*_, and total activity, *T*_*i*_ = ∑_*j*_ *a*_*ij*_. Reported proteome summaries are arithmetic means: 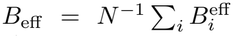 , *q*_max_ = *N* ^−1^ ∑_*i*_ *q*_*i*,max_, 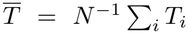, and 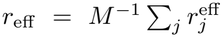. Weakest-reaction performance is

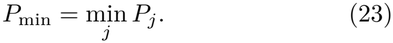

A change Δ*X* denotes the endpoint value minus the ancestral value for the same run. Unless stated otherwise, endpoints are means of five replicates, and error bars and reported ± values denote one sample standard deviation. Faint points show individual replicates. The reference trajectory is a single simulation rather than a replicate mean.

## III. ANALYTICAL EXPECTATIONS

The stochastic simulations are not intended to admit a closed-form evolutionary trajectory. However, several consequences of the phenotype-level fitness function provide mechanistic expectations against which the numerical results can be interpreted.

### A. Redundancy lowers the selective cost of activity loss

Suppose protein *i* contributes

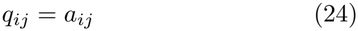

to reaction *j*, and a mutation completely removes that contribution. Then

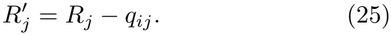

If no other reaction changes, the exact relative fitness is

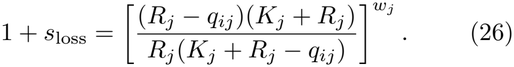

If *r* proteins contribute equally while total reaction capacity *R*_*j*_ is held fixed,

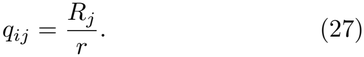

For a sufficiently small contribution loss,

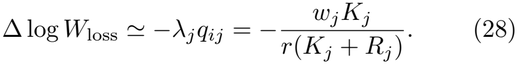

Hence

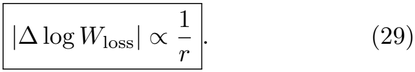

Thus, at fixed total reaction capacity, distributed redundancy lowers the selective cost of removing one contributor. This is a local buffering result rather than a prediction that the full evolutionary endpoint must scale as 1*/r*.

### B. Selection on loss-containing pleiotropic mutations

For a sufficiently small pleiotropic mutation,

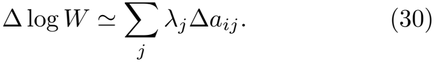

A mutation containing both gains and losses is favoured to first order when

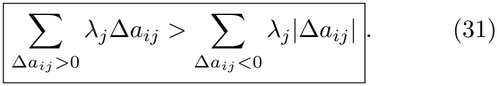

For one gain and one loss this reduces to

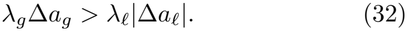

Redundancy therefore does not make biochemical loss intrinsically beneficial. Rather, by lowering the marginal selective value *λ*_*j*_ of activity in well-supplied reactions, it can reduce the organismal cost associated with some losses sufficiently that a pleiotropic mutation containing those losses remains advantageous.

The simulations themselves use the exact finite selection coefficient; the expressions above provide the small-effect mechanistic interpretation.

### C. Dimensionless structural incompatibility

For a loop supporting *k* activities of equal magnitude *a*,

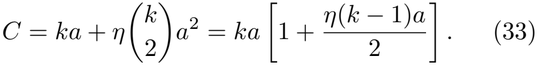

The ratio of the pairwise incompatibility contribution to the linear cost is therefore

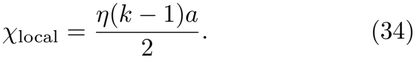

The parameter used in the simulations evaluates this ratio at the ancestral reference state,

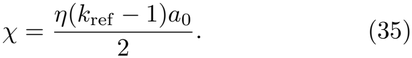

Thus

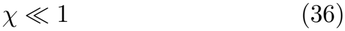

corresponds to weak pairwise incompatibility relative to the linear loop cost,

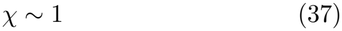

to comparable pairwise and linear contributions, and

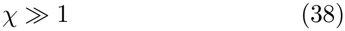

to a feasibility constraint dominated by simultaneous multifunctionality.

This dimensional analysis does not imply that specialisation must increase monotonically with *χ*; the evolutionary response also depends on how structural feasibility reshapes the accessible mutational neighbourhood.

## IV. RESULTS

### A. Greater redundancy permits stronger specialisation at matched evolutionary exposure

The ancestral proteins were broad but not perfectly even, with a mean effective breadth of approximately 13.5 compared with a nominal support of 15 reactions. After an evolutionary exposure of *E* = 5 fixed substitutions per protein, effective breadth decreased across every parameter combination examined (Fig. 1A). This was accompanied by an increase in the fraction of each protein’s activity carried by its strongest function (Fig. 1B), showing that the change in breadth reflected a genuine concentration of catalytic activity.

**FIG. 1.**
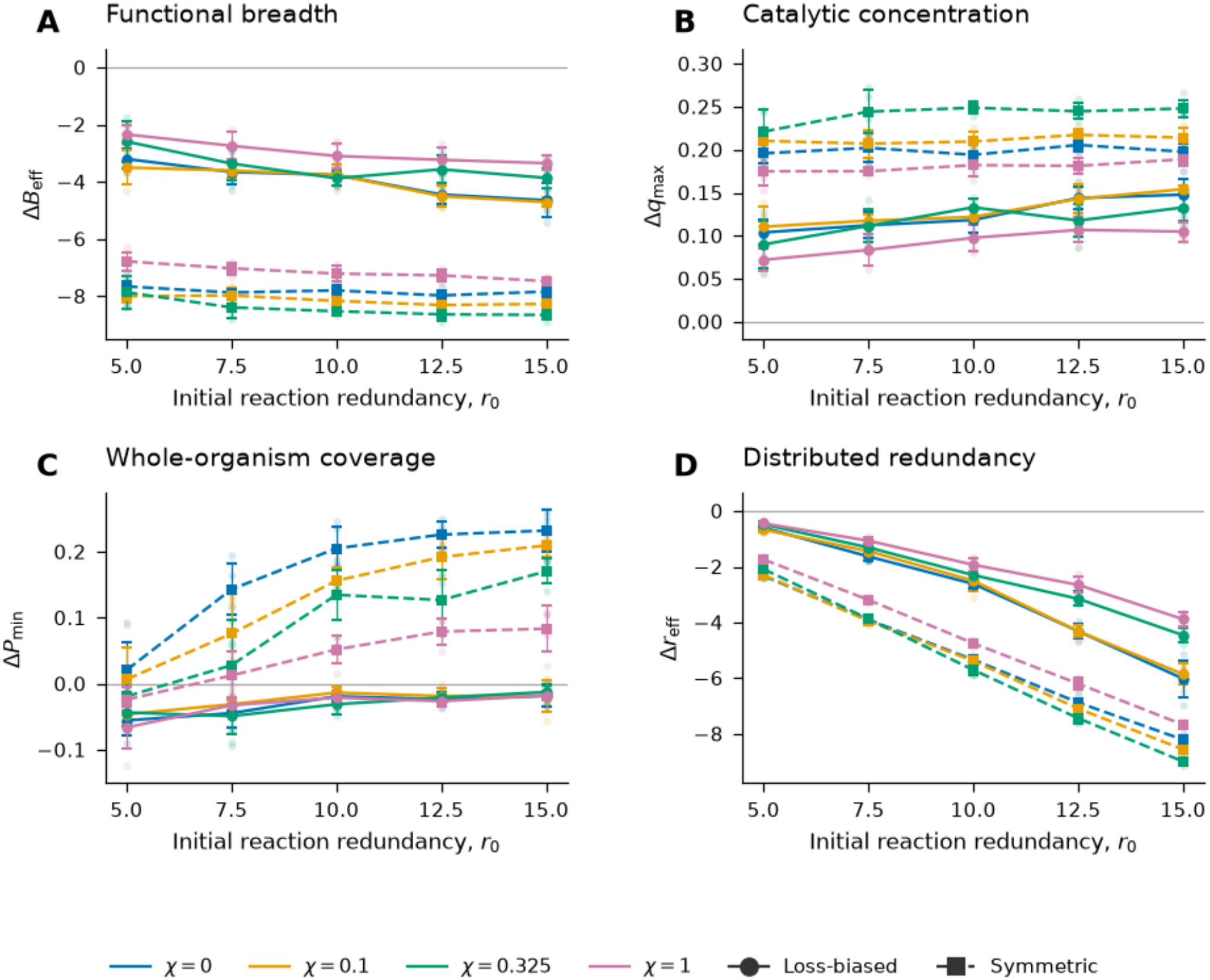
Specialisation and reaction performance at matched evolutionary exposure. All conditions are compared after *E* = *n*_fix_*/N* = 5 fixed substitutions per protein. Colours denote structural incompatibility *χ*; solid circles show the loss-biased mutation kernel and dashed squares the symmetric kernel. Points show replicate means and error bars denote one sample standard deviation. (A) Change in mean effective protein breadth, Δ*B*_eff_. (B) Change in mean strongest-function share, Δ*q*_max_. (C) Change in weakest-reaction performance, Δ*P*_min_. (D) Change in mean effective reaction redundancy, Δ*r*_eff_. Greater initial redundancy generally produces stronger protein-level specialisation under the loss-biased kernel while reducing the loss of weakest-reaction performance.

Under the loss-biased mutation kernel, increasing initial reaction redundancy generally produced stronger specialisation. Across the tested structural-incompatibility values, higher-*r*_0_ proteomes typically showed a larger reduction in effective breadth and a larger increase in strongest-function share at the same evolutionary exposure. The effect was clearest for weak to intermediate structural incompatibility, while the response was weaker at *χ* = 1. This dependence on redundancy is also visible throughout the evolutionary trajectories (Fig. 2A–D). At the beginning of evolution the loss-biased trajectories are similar, but proteins in the higher-redundancy systems progressively reach lower effective breadth at the same number of fixed substitutions per protein. The separation therefore develops during evolution rather than arising from differences in the ancestral states.

**FIG. 2.**
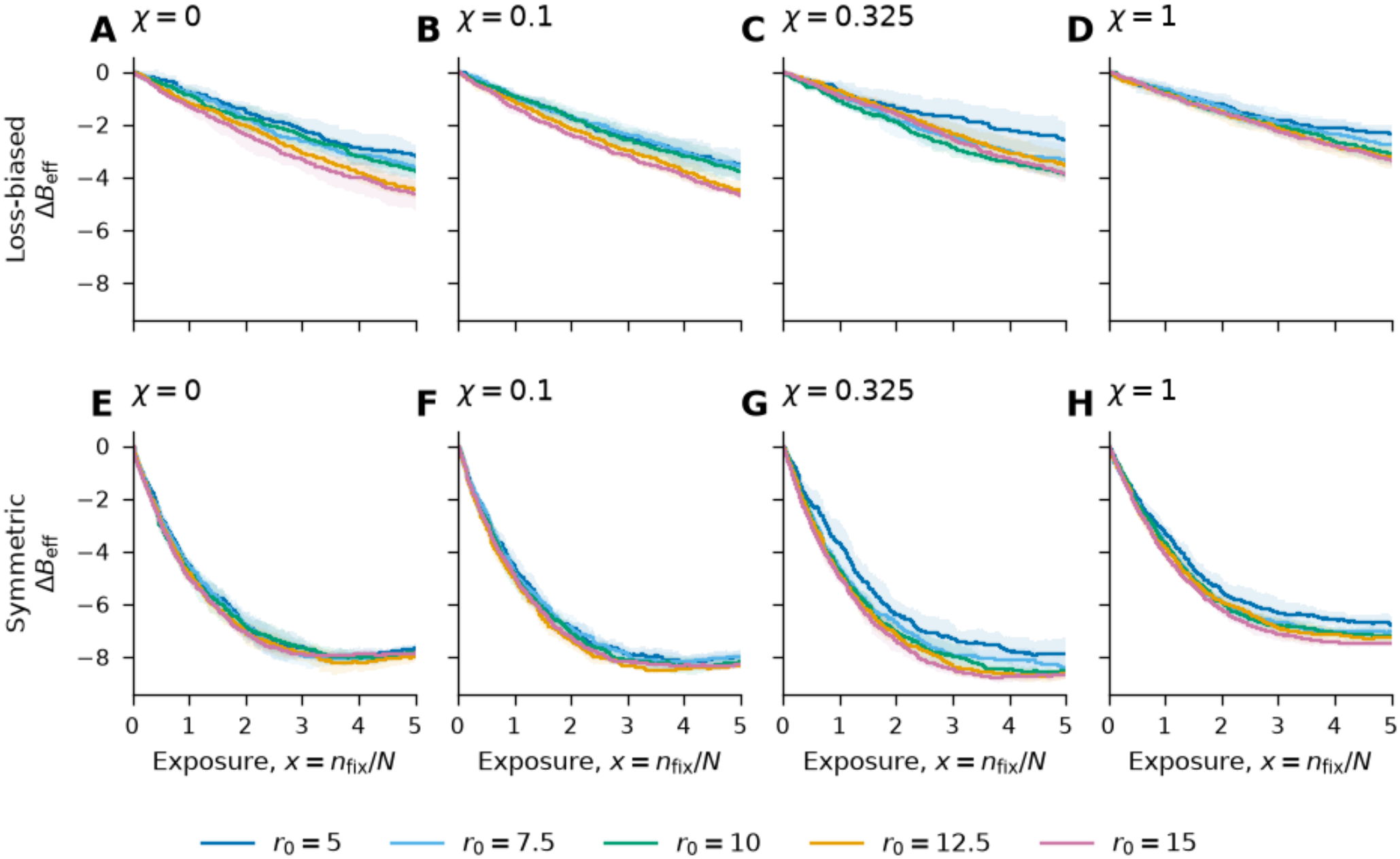
Evolution of effective protein breadth at matched exposure. Change in mean effective breadth is shown as a function of evolutionary exposure, *E* = *n*_fix_*/N* , for each initial redundancy *r*_0_. Panels (A–D) show the loss-biased mutation kernel and panels (E–H) the symmetric kernel, with structural incompatibility increasing from *χ* = 0 to *χ* = 1. Lines show replicate means and shaded regions show variation among replicates. Under loss-biased mutation, higher-redundancy systems generally undergo greater narrowing at the same evolutionary exposure. Symmetric mutation produces substantially stronger and more rapid breadth reduction, with trajectories tending to approach similar lower-breadth states.

### B. Specialisation remains compatible with organismal reaction coverage

The additional protein-level specialisation observed at higher redundancy did not produce a corresponding deterioration in organismal reaction performance. Under loss-biased mutation, weakest-reaction performance declined modestly at low initial redundancy but was increasingly preserved as *r*_0_ increased (Fig. 1C). At the highest redundancies examined, the mean change in *P*_min_ was close to zero for several structural-incompatibility conditions.

This produces the combination expected from distributed functional buffering: individual protein profiles become more concentrated while the least well supplied reaction remains comparatively stable. The result is consistent with the analytical expectation that the selective cost of removing one contribution decreases as the number of contributors to a reaction increases.

Effective reaction redundancy nevertheless decreased during evolution (Fig. 1D). The decline became larger in absolute terms as initial redundancy increased, indicating that the proteome did not simply retain an increasingly diffuse set of equivalent contributors. Instead, protein-level specialisation was accompanied by a reorganisation of how catalytic activity was distributed among the proteins supplying each reaction.

### C. Mutation regime and structural incompatibility shape the extent of specialisation

The symmetric mutation kernel produced substantially stronger specialisation than the loss-biased kernel across the parameter grid (Fig. 1A,B). This difference is also apparent in the exposure trajectories (Fig. 2). Under symmetric mutation, effective breadth falls rapidly during the first few substitutions per protein and then approaches a lower-breadth regime. In contrast, the loss-biased trajectories decline more gradually and generally remain in motion at the end of the simulated exposure.

The effect of initial redundancy was consequently less pronounced under symmetric mutation. Strong narrowing occurred even at low *r*_0_, and trajectories with different redundancies often approached similar effective breadth at later exposure (Fig. 2E–H). A bias towards biochemical loss is therefore not required for specialisation in this model; the availability of positive changes and redistribution of catalytic activity can support even stronger narrowing.

Structural incompatibility modified the magnitude of the response but did not act as a simple monotonic driver. The strongest constraint examined, *χ* = 1, did not consistently produce the greatest specialisation. This is consistent with the structural term acting as a constraint on accessible states rather than as a direct selective reward for narrow activity profiles.

### D. Representative dynamics and catalytic output

A representative loss-biased trajectory shows how the different components of specialisation change together (Fig. 3B,C). Effective breadth declines throughout the trajectory while the strongest-function share rises. In contrast, weakest-reaction performance remains comparatively close to its ancestral value. Specialisation in this run therefore reflects increasing concentration of activity within individual proteins rather than a comparable loss of organismal reaction coverage.

**FIG. 3.**
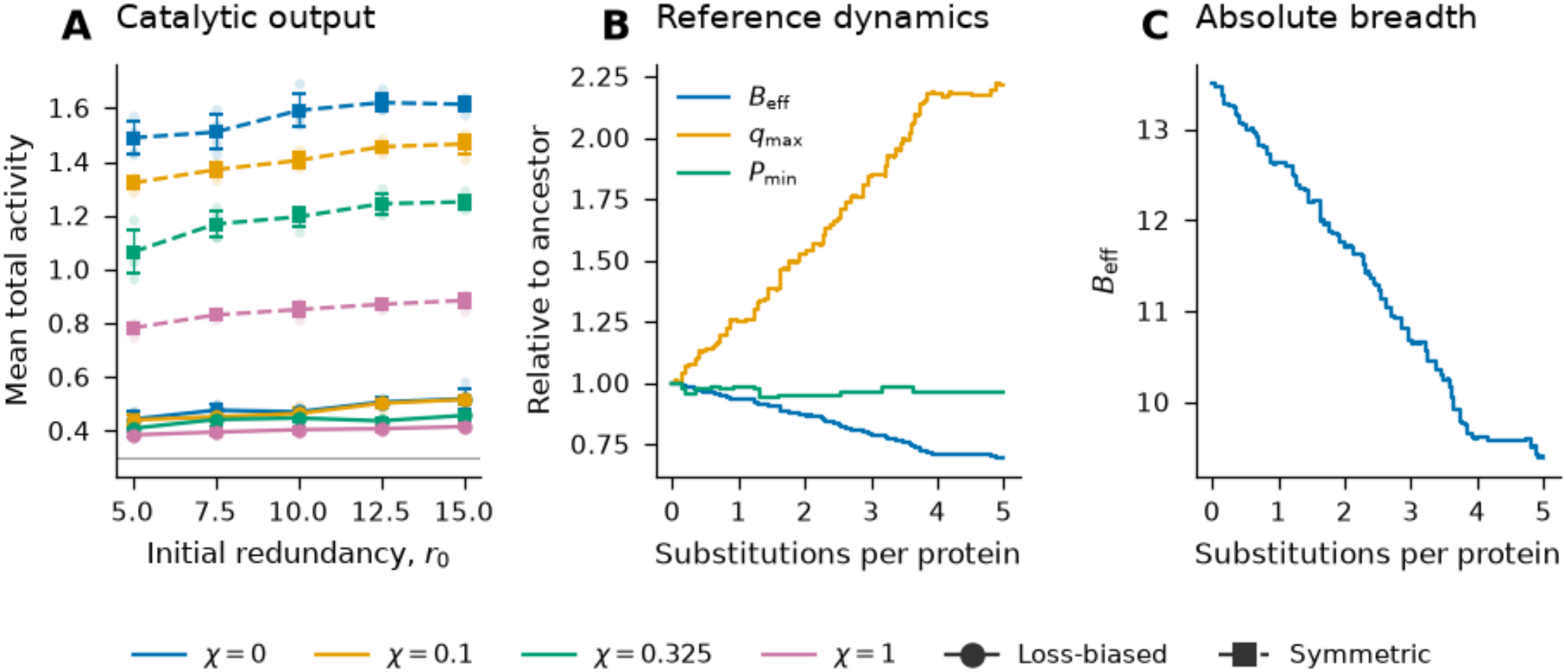
Catalytic output and representative evolutionary dynamics. (A) Mean total catalytic activity per protein at *E* = 5 across the redundancy sweep. Colours denote structural incompatibility *χ*; solid circles show the loss-biased mutation kernel and dashed squares the symmetric kernel. Error bars denote one sample standard deviation across replicates. (B) Representative trajectory for the reference loss-biased condition (*r*_0_ = 12.5, *χ* = 0.325), showing effective breadth *B*_eff_, strongest-function share *q*_max_, and weakest-reaction performance *P*_min_ relative to their ancestral values as a function of fixed substitutions per protein. (C) Absolute effective breadth for the same trajectory. Effective breadth continues to decline at the end of the simulated exposure, so *E* = 5 is treated as a common comparison point rather than an inferred evolutionary equilibrium.

The absolute breadth trajectory is still declining when the simulation reaches *E* = 5 (Fig. 3C). We therefore use *E* = 5 as a matched observation point across redundancy conditions, not as an estimate of the eventual evolutionary endpoint. This behaviour is also consistent with the wider set of loss-biased trajectories in Fig. 2, many of which remain in motion at the end of the sampled exposure.

Mean total catalytic activity also increases during evolution (Fig. 3A), particularly under the symmetric mutation kernel. Because total activity is not conserved by the model, this increase is not itself evidence for specialisation or improved efficiency. The relevant signatures of specialisation are instead the decrease in effective breadth and the increase in strongest-function share.

## V. DISCUSSION

The main result is that distributed functional overlap can allow individual proteins to specialise while organismal reaction performance remains buffered. At matched evolutionary exposure, higher initial redundancy generally produces a larger reduction in effective breadth under loss-biased and unbiased mutation kernel, while the least well supplied reaction is better preserved. This is consistent with the analytical result above: when several proteins contribute to the same reaction, the selective cost of reducing one contribution decreases approximately as 1*/r*. Redundancy therefore does not make loss beneficial by itself, but it can make some losses cheap enough that pleiotropic mutations with gains elsewhere remain favourable. This kind of context dependence is consistent with systems-level views of enzyme evolution [12] and with multicopy-suppression experiments, where weak or noncognate activities can sometimes compensate for deficiencies elsewhere in metabolism [13].

The mutation-kernel comparison also shows that loss bias is not required for specialisation. The symmetric kernel produces substantially stronger narrowing and often approaches a lower-breadth regime within the exposure studied here. This is not a clean test of mutation sign alone, because the symmetric kernel also changes the magnitude distribution and makes larger positive changes more available. Even so, the result suggests that specialisation depends on opportunities to amplify and redistribute activity, not only on the removal of weak secondary functions. That is broadly consistent with experimental work showing that promiscuous activities can improve substantially while the original activity is retained, although trade-offs can emerge later in optimisation [10, 11]. Structural incompatibility shows a similarly non-monotonic effect: increasing *χ* makes simultaneous activities harder to maintain, but also removes accessible mutational states, so stronger constraint does not automatically produce stronger specialisation.

The endpoint at *E* = 5 should not be read as an evolutionary equilibrium. Several loss-biased trajectories are still declining in breadth at the end of the sampled exposure, including the representative trajectory in Fig. 3. We therefore use *E* = 5 as a common observation point rather than an estimate of the eventual degree of specialisation. Total catalytic activity is also not conserved and generally increases during evolution, particularly under the symmetric kernel. For that reason, the evidence for specialisation comes from redistribution of activity within proteins: effective breadth decreases while strongest-function share increases. The increase in total activity should not be interpreted as an efficiency gain from reallocating a fixed biochemical resource.

Several features of real enzyme evolution are deliberately absent from the model. The activities *a*_*ij*_ are phenomenological rather than measured kinetic constants, protein abundances and reaction weights are equal, and the set of organismal reactions is fixed. The structural constraint is a coarse-grained description of local incompatibility and does not explicitly model sequence, folding, substrate binding, or metabolic flux. The mutation kernels are likewise phenomenological rather than fitted to an empirical distribution of biochemical effects. Finally, the origin–fixation framework describes sequential fixation using the standard diffusion approximation [14] and does not include competition among simultaneously segregating variants. The results should therefore be read as a mechanistic result within a minimal model, rather than as a quantitative reconstruction of a particular proteome.

Gene duplication remains one of the major routes by which enzymes diversify and specialise [3–8]. The present model does not compete with that history, and the over-lapping proteome used as its starting point could itself have arisen through earlier duplication, recruitment, or other evolutionary processes. The narrower point is that, once several existing proteins already contribute to the same reactions, another duplication event is not required before their catalytic roles can begin to separate. Distributed overlap reduces the organismal consequence of changing one contributor, and in these simulations that buffering allows greater protein-level specialisation while the collective reaction set remains supported.

## ACKNOWLEDGMENTS

OpenAI’s ChatGPT assisted with literature searching and synthesis, manuscript drafting and editing, and LATEX preparation, following author-specified constraints to preserve the model and numerical results. The authors take full responsibility of all scientific arguments, model assumptions, numerical results, interpretations, and final manuscript content.

## DATA AND CODE AVAILABILITY

The simulation code, parameter-sweep outputs, and plotting scripts used to generate the results and figures in this study are publicly available in the accompanying GitHub repository [15].

## References

[1] P. J. O’Brien and D. Herschlag, Catalytic promiscuity and the evolution of new enzymatic activities, Chem. Biol. 6, R91 (1999).

[2] O. Khersonsky and D. S. Tawfik, Enzyme promiscuity: A mechanistic and evolutionary perspective, Annu. Rev. Biochem. 79, 471 (2010).

[3] K. Voordeckers, C. A. Brown, K. Vanneste, E. van der Zande, A. Voet, S. Maere, and K. J. Verstrepen, Reconstruction of ancestral metabolic enzymes reveals molecular mechanisms underlying evolutionary innovation through gene duplication, PLoS Biol. 10, e1001446 (2012).

[4] S. Ohno, Evolution by Gene Duplication (Springer, Berlin, Heidelberg, 1970).

[5] A. Force, M. Lynch, F. B. Pickett, A. Amores, Y.-l. Yan, and J. Postlethwait, Preservation of duplicate genes by complementary, degenerative mutations, Genetics 151, 1531 (1999).

[6] S. D. Copley, Evolution of new enzymes by gene duplication and divergence, FEBS J. 287, 1262 (2020).

[7] U. Bergthorsson, D. I. Andersson, and J. R. Roth, Ohno’s dilemma: Evolution of new genes under continuous selection, Proc. Natl. Acad. Sci. U.S.A. 104, 17004 (2007).

[8] J. Näsvall, L. Sun, J. R. Roth, and D. I. Andersson, Real-time evolution of new genes by innovation, amplification, and divergence, Science 338, 384 (2012).

[9] R. A. Jensen, Enzyme recruitment in evolution of new function, Annu. Rev. Microbiol. 30, 409 (1976).

[10] A. Aharoni, L. Gaidukov, O. Khersonsky, S. M. Gould, C. Roodveldt, and D. S. Tawfik, The ‘evolvability’ of promiscuous protein functions, Nat. Genet. 37, 73 (2005).

[11] N. Tokuriki, C. J. Jackson, L. Afriat-Jurnou, K. T. Wyganowski, R. Tang, and D. S. Tawfik, Diminishing returns and tradeoffs constrain the laboratory optimization of an enzyme, Nat. Commun. 3, 1257 (2012).

[12] S. D. Copley, Toward a systems biology perspective on enzyme evolution, J. Biol. Chem. 287, 3 (2012).

[13] W. M. Patrick, E. M. Quandt, D. B. Swartzlander, and I Matsumura, Multicopy suppression underpins metabolic evolvability, Mol. Biol. Evol. 24, 2716 (2007).

[14] M. Kimura, On the probability of fixation of mutant genes in a population, Genetics 47, 713 (1962).

[15] A. Duygu and H. Moinudeen, Code and data for “a minimal model of functional redundancy as a route to enzyme specialisation”, https://github.com/haziqmoinudeen-git/alp-duygu-functional-redudancy (2026), gitHub repository.

